# Convective forces increase CXCR4-dependent glioblastoma cell invasion in GL261 murine model

**DOI:** 10.1101/451286

**Authors:** R. Chase Cornelison, Caroline E. Brennan, Kathryn M. Kingsmore, Jennifer M. Munson

**Affiliations:** Department of Biomedical Engineering and Mechanics, Virginia Polytechnic Institute and State University, Blacksburg, VA 24061; Department of Biomedical Engineering, University of Virginia, Charlottesville, VA 22908

**Keywords:** Glioblastoma,, invasion,, interstitial fluid flow,, convection enhanced delivery,, CXCR4,, CXCL12

## Abstract

Glioblastoma is the most common and malignant form of brain cancer. Its invasive nature limits treatment efficacy and promotes inevitable recurrence. Previous *in vitro* studies have shown that interstitial fluid flow, a factor characteristically increased in cancer, increases glioma cell invasion via CXCR4-CXCL12. It is currently unknown if these effects translate *in vivo.* Using the therapeutic technique of convection enhanced delivery (CED), we tested if convective flow alters glioma invasion *in vivo* using the syngeneic GL261 mouse model of glioblastoma. We first confirmed that GL261 invasion *in vitro* increased under flow in a CXCR4-CXCL12 dependent manner. Additionally, approximately 65.4% and 6.59% of GL261 express CXCR4 and CXCL12 *in vivo,* respectively, with 3.38% expressing both. Inducing convective flow within implanted tumors indeed increased glioma cell invasion over untreated controls, and administering CXCR4 antagonist AMD3100 (5 mg/kg) effectively eliminated this response. Therefore, glioma invasion is in fact stimulated by convective flow *in vivo* through CXCR4. We also analyzed patient samples to show that expression of CXCR4 and CXCL12 increase in patients following therapy. These results suggesting that targeting flow-stimulated invasion may prove beneficial as a second line of therapy, particularly in patients chosen to receive convection enhanced drug delivery.

## Introduction

Glioblastoma (GBM) is the most aggressive form of brain cancer and is characterized by invasion into the surrounding brain or parenchyma^1,2^. This invasiveness causes diffuse borders between the tumor and parenchyma, preventing effective resection of all malignant cells. Additionally, because tumor cells that have invaded into the surrounding healthy tissue are increasingly resistant to radiation and chemotherapy, GBM always recurs^3,4^. Therefore, understanding and targeting molecules that regulate glioma cell invasion has therapeutic implications in the treatment of GBM. One signaling axis known to regulate GBM invasion is the CXCR4-CXCL12 pathway. While a potent driver of GBM invasion in static conditions, CXCR4-and CXCL12-mediated invasion in GBM can be enhanced by interstitial fluid flow through a mechanism known as autologous chemotaxis^5-7^. Interstitial flow is the movement of fluid from the vasculature throughout the interstitial tissue space toward draining lymphatics or clearance pathways. This process normally acts to maintain tissue homeostasis, but the leaky nascent vasculature and increased waste production in solid cancers can dramatically increase interstitial pressure and, in turn, interstitial flow^1,8^.

We previously showed that rat and human GBM cell lines respond to flow *in vitro* by increasing invasion^6,7^. Furthermore, regions of high flow (identified by arterial extravasation of Evans blue) correlated with regions of invasion for cell lines as well as patient-derived glioma stem cells^6,7^. Flow-stimulated invasion was mitigated by both blocking the receptor CXCR4 as well as saturating the ligand CXCL12, suggesting this chemokine-receptor pathway plays a key role in glioma cell flow response. It remains unknown, however, if interstitial flow directly stimulated cancer cell invasion *in vivo* and if CXCR4 signaling was similarly implicated. Answering these questions requires a technique to induced convective forces within the tumor *in situ* at a time when heightened interstitial flow may not be fully established on its own.

Convection enhanced delivery (CED) is an experimental technique used in the clinic to overcome high intra-tumoral pressure and increase drug distribution via local infusion^9,10^. A blunt needle is placed into the center of the tumor, and a drug-laden solution is infused to drive drug transport. In essence, CED drives convective flow through the interstitial spaces in the tumor, mimicking interstitial fluid flow. We used CED in a murine model of GBM to test the hypothesis that convective flow directly stimulates cancer cell invasion *in vivo* and examine the dependence of this response on CXCR4 signaling.

## Results

### GL261 exhibit flow-stimulated invasion in vitro in a CXCR4-dependent manner

Prior to *in vivo* assessment, the flow response of GL261 cells was examined *in vitro* using a 3D tissue culture insert model (**Fig. 1A**)^6^. Under static conditions, 0.1-0.2% of GL261 invaded beyond the semipermeable membrane (**Fig. 1B**). The addition of gravity-driven flow significantly increased the percent of cells invading by approximately 1.6 fold (t(4)=5.931, n=5, p<0.01). This flow-stimulated increase in invasion could be mitigated by blocking CXCR4 using 10 μM AMD3100, a small molecule inhibitor of CXCR4 (t(4)=2.722, n=5, p>0.1). Similar results were observed for saturating the cultures (in the gel and both sides of the transwell) with 100 nM CXCL12. Ligand saturation significantly decreased the effects of flow (t(4)=3.545, n=5, p<0.05) (**Fig. 1C**), returning invasion to untreated levels (t(3)=2.293, n=4, p>0.1). Hence, the flow response of GL261 aligns with the previously proposed mechanism of CXCR4-CXCL12 autologous chemotaxis^1^.

**Figure 1:**
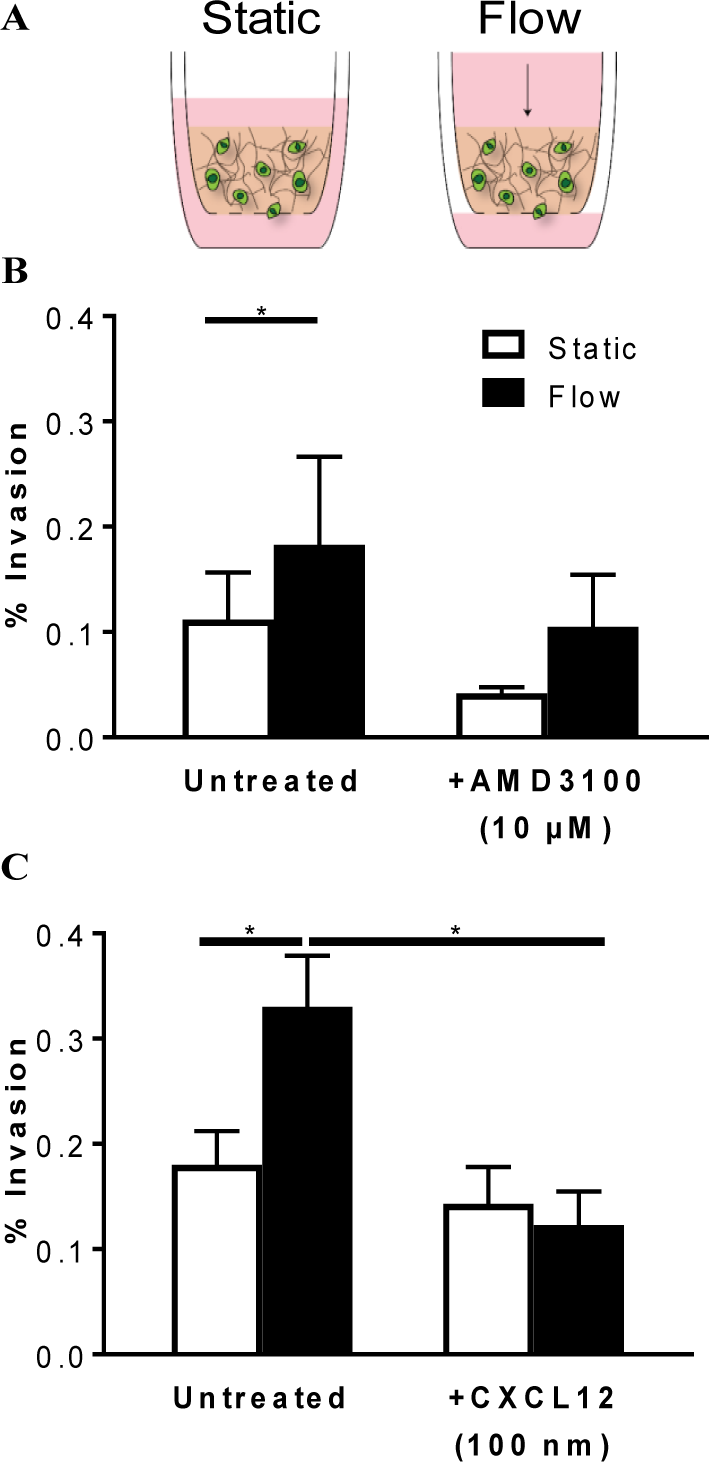
Interstitial flow increases GL261 invasion in a CXCR4-CXCL12 dependent manner. A) Schematic representation of tissue culture insert setup for static and flow experimental conditions. B) Percent invasion of GL261 in static and flow conditions with and without addition of 10 μM AMD3100 (n=5, *p=0.01). C) Percent GL261 invasion in static and flow conditions with and without addition of 100 nM CXCL12 (n=4, *p=0.01). Bars show standard error.

### CXCR4^+^ and CXCR4+CXCL12^+^ populations are enriched within in vivo tumor samples

Because the significance of targeting autologous chemotaxis and flow-stimulated invasion may be influenced by expression levels, we used flow cytometry to characterize GL261 expression of CXCR4 and CXCL12 in different environments. The dimensionality of culture significantly impacted receptor and ligand expression. In 2D, few cells expressed the receptor, ligand, or both (**Fig. 2**). Embedding the cells in 3D hydrogels significantly increased the number of CXCR4^+^ cells to 8.13 ± 1.71% compared to 1.83 ± 0.25% in 2D culture (t(3)=3.389, n=4, p<0.05) (**Fig. 2A**). Similar effects were observed on the CXCL12 population (t(3)=4.14, n=4, p<0.05) (**Fig. 2B**). While there was no difference in the percentage of CXCR4^+^CXCL12^+^ cells between 2D and 3D *in vitro* culture (**Fig. 2C**), this double positive population increased from 1.66 ± 0.72% in 3D to 3.38 ± 0.49% of total cells *in vivo* (t(8)=2.767, n=6 *in vivo* and n=4 *in vitro,* p<0.05 compared to 3D). These effects were further amplified for CXCR4 single expression, dramatically increasing from 8.13 ± 1.71% in 3D to 65.4 ± 5.19% *in vivo* (t(8)=8.653, n=6 *in vivo* and n=4 *in vitro,* p<0.0001 compared to 3D). Expression of CXCL12 *in vivo* was similar to that in 3D culture. Given the role of this receptor/ligand pair on flow response, an enrichment in CXCR4^+^ and CXCR4+CXCL12+ populations may increase the potential for flow-stimulated invasion *in vivo*.

**Figure 2:**
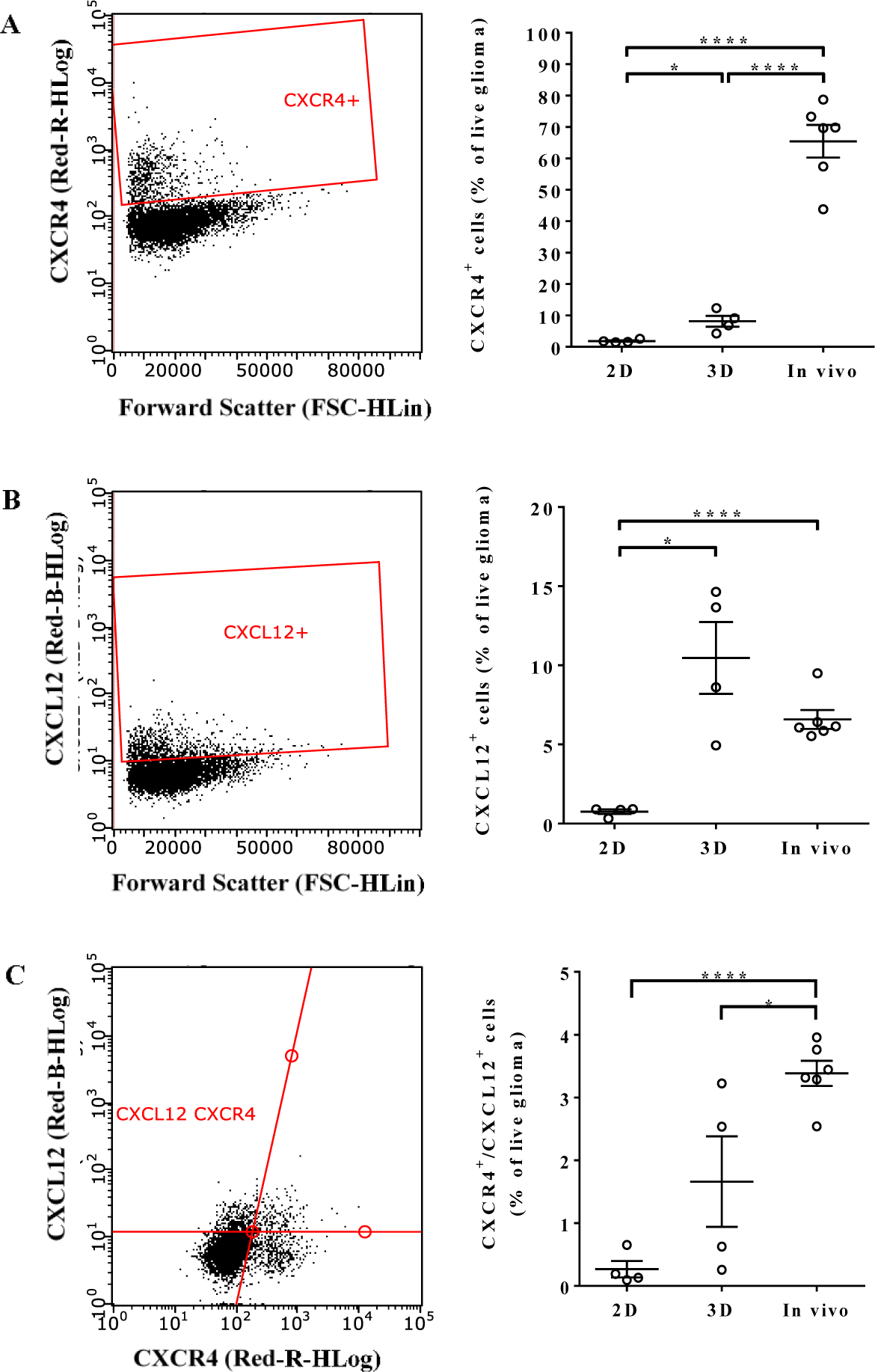
Population-level expression of CXCR4 and CXCL12 in GL261 depends on growth conditions. Flow cytometry was used to determine the percent of CXCR4+, CXCL12+, and double positive GL261 in 2D, 3D, and *in vivo* environments. Representative plots gated on live glioma cells are shown in the left column for (A) CXCR4, (B) CXCL12, and (C) double positive populations. Correlating quantifications are shown on the right. *p<0.05, ****p<0.0001. Bars

### Glioma invasion in vivo is enhanced by convective flow

We examined the effects of convective forces on glioma cell invasion *in vivo*using the therapeutic technique of convection enhanced delivery (CED). A cartoon of the process is shown in **Figure 3A**, and an experimental timeline in **Figure 3B**. First, magnetic resonance imaging was used to verify the ability to induce fluid convection using CED. A gadolinium contrast agent conjugated to albumin (Galbumin, 25 mg/mL) was infused into the tumors at day 7 at a rate of 1 μL/min. Immediately following CED, the mice were transferred to a 7 Tesla MRI machine to visualize changes in galbumin distribution over time. Five representative slices are shown for one mouse. T2-weighted images were used to identify the location of the tumors (**Suppl. Fig. 2A**). Using T1-weighted imaging, the signal intensity of intra-tumoral galbumin was observed to change over a 30 minute period, indicative of contrast agent flux (**Suppl. Fig. 2B**).

**Figure 3:**
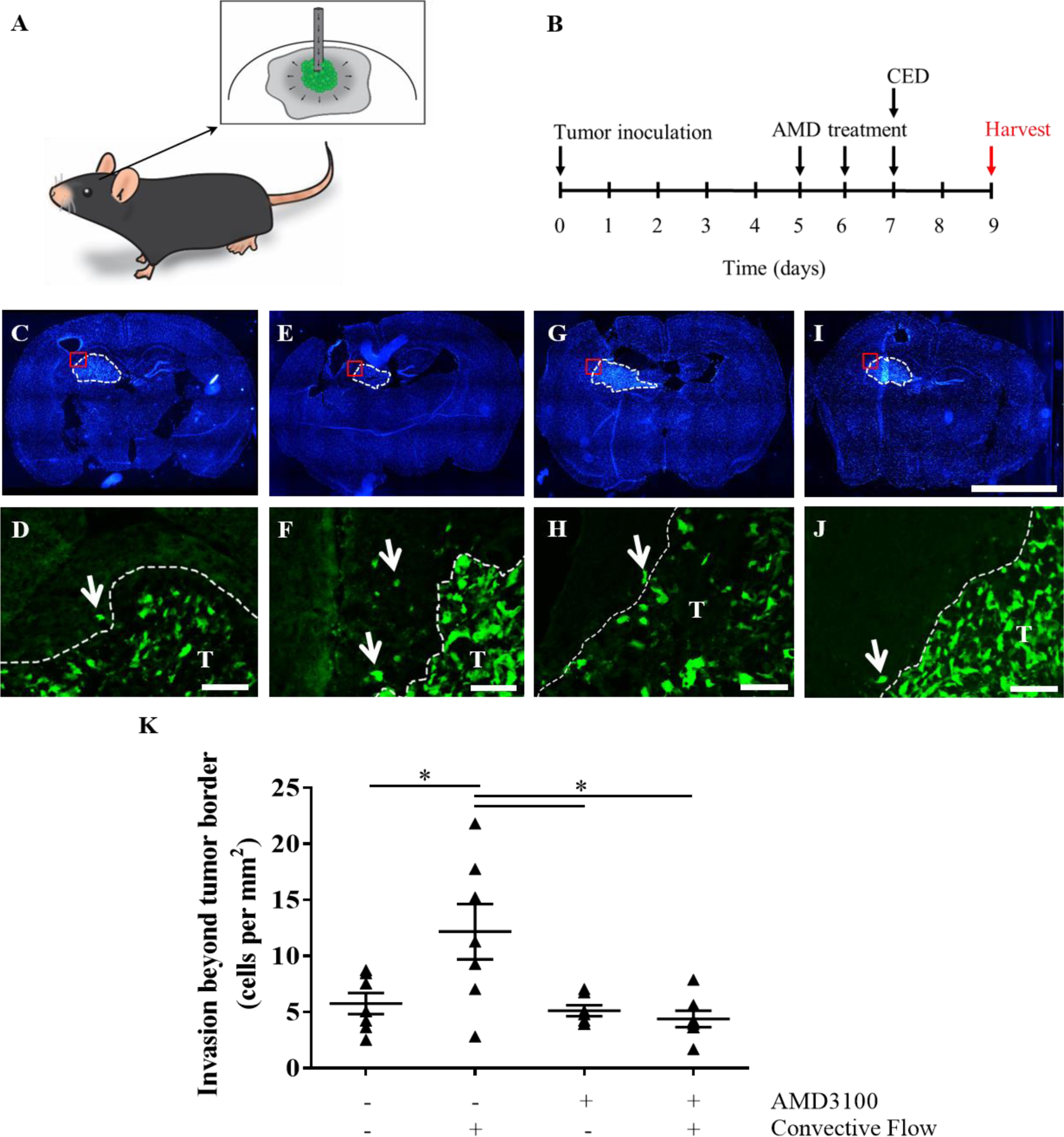
Interstitial flow increases murine glioma cell invasion *in vivo* in a CXCR4-dependent manner. (A) Schematic of intratumoral convection enhanced delivery. (B) Experimental timeline. (C-J) Representative fluorescence images of *in vivo* glioma invasion for (C-D) untreated controls, (E-F) CED alone group, (G-H) AMD alone group, and (I-J) +CED/+AMD group. Top: Full brain slice scans with nuclei labeled with DAPI (blue), with tumor defined by white dotted line. Scale bar= 1 mm. Bottom: GFP-labeled GL261 tumor cells at the border location depicted above (red boxes). Scale bar=100μm. (K) Quantification of tumor cells beyond the tumor border averaged per mouse from five locations in three sections through tumors. Bars show standard error.

Following verification that CED does induce convective flow, a second cohort of mice was used to examine invasion. Convective flow was again induced seven days after tumor inoculation at 1 μL/min, and invasion was assessed two days later using immunohistochemistry. Representative images are shown in **Figures 3C-J**, with invasion quantification summarized in **Figure 3K**. Untreated (static) tumors had approximately 5.75 ± 0.938 cells/mm^2^ invaded beyond the tumor border into the surrounding tissue. Following CED, the number of invading cells significantly increased to 12.2 ± 2.4/mm^2^ (**Fig. 3E-F, K**) (t(12)=2.433, n=7, p<0.05). This greater than 2-fold increase to invasion *in vivo* was even more pronounced than the *in vitro* results (increased approximately 1.5-fold under flow).

### Effects of flow in vivo are mediated through CXCR4

Given the ability of CXCR4 antagonism to reduce flow-stimulated invasion *in vitro,* we also examined the effects of administering the CXCR4 antagonist AMD3100 (5 mg/kg) systemically with and without CED^11^. This drug has been delivered to *in vivo* glioma models previously and shows some clinical potential as a secondary therapy^12,14^. In the absence of convective flow (**Fig. 3G-H, K**), AMD3100 did not significantly alter glioma cell invasion compared to untreated controls at 5.12 ± 0.490/mm^2^(t(12)=0.6008, n=7, p>0.1). However, applying CED in mice treated with AMD3100 (**Fig. 3I-J, K**) significantly reduced the effects of flow on invasion compared to CED alone (t(12)=3.026, n=7, p<0.05). This treatment regimen effectively maintained the number of cells invading beyond the tumor border to 4.38 ± 0.731 cells/mm^2^, not significantly different from that of untreated, static controls. Hence, dosing with AMD3100 prior to convection is able to mitigate flow-stimulated increases to glioma cell invasion. This decrease in flow-stimulated invasion with AMD3100 treatment was associated with a decrease in CXCR4 phosphorylation, an indicator of receptor stimulation and signaling^6^. Untreated tumors exhibited moderate immunoreactivity for phosphorylated CXCR4 (**Fig. 4A**), indicating that this signaling pathway is basally active within GL261 tumors *in vivo.* Consistent with prior *in vitro* results, applying flow via CED markedly increased pCXCR4 immunoreactivity *in vivo* (**Fig. 4B**). Administering AMD3100 prior to CED effectively attenuated increased pCXCR4 staining, observably decreasing immunoreactivity below that of untreated controls (**Fig. 4C**). No qualitative differences were observed in total CXCR4 expression (data not shown). Hence, interstitial flow is indeed able to stimulate invasion of glioma cells *in vivo* mediated at least in part through CXCR4 signaling.

**Figure 4:**
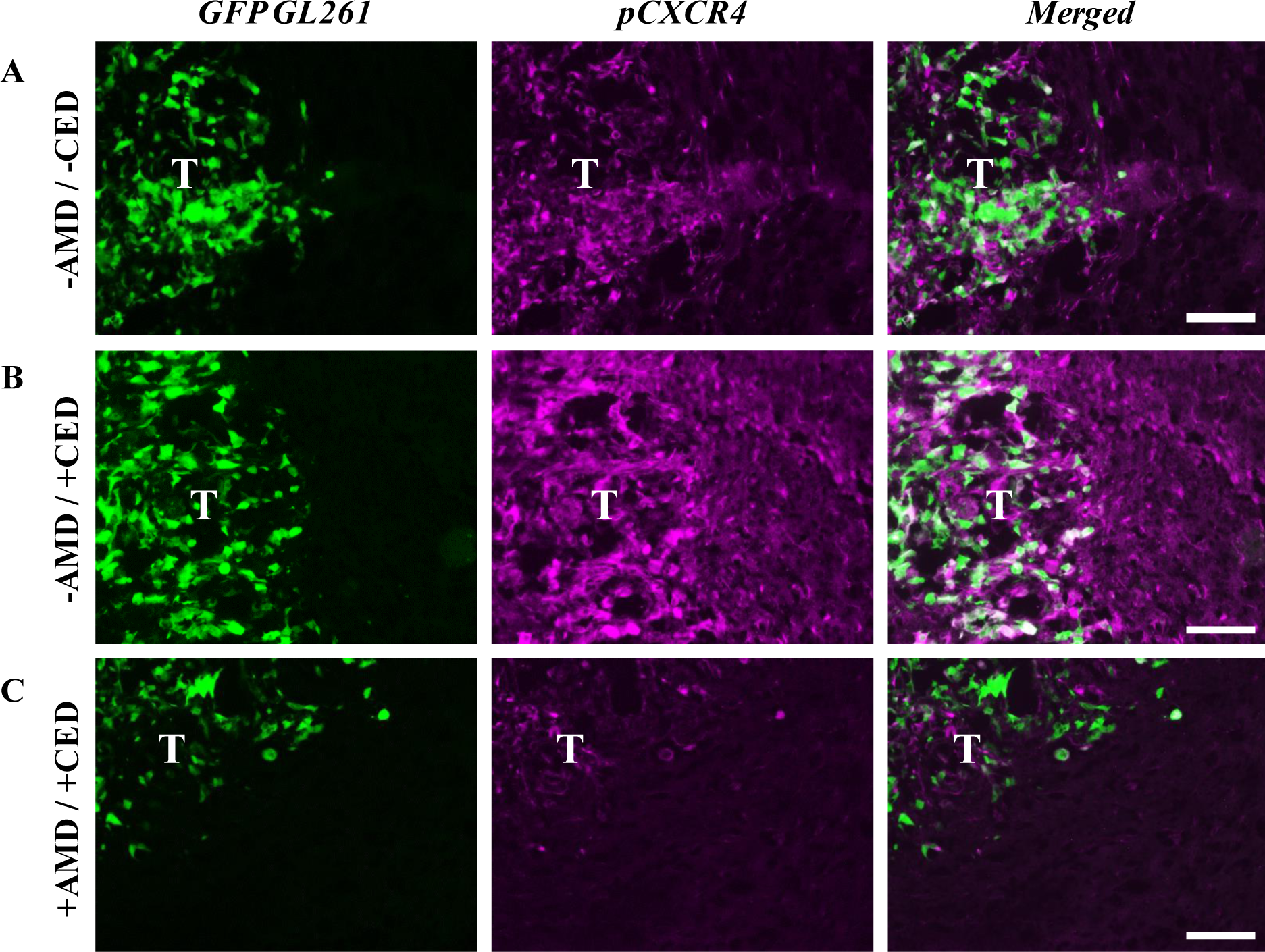
Treatment of GL261 with AMD3100 decreases convection-driven increases in pCXCR4. Representative fluorescence images at GL261 tumor (T) borders of GFP-GL261 (green) and pCXCR4 (magenta) in (A) untreated animals, (B) animals receiving only CED, and (C) animals dosed with AMD3100 for two days prior to CED. Scale bars = 100 μm.

### Standard of care therapy may increase CXCR4 and CXCL12 expression in patients

Convection enhanced delivery is experimentally used in the clinic to deliver a secondary therapy, meaning it is implemented after the standard of care radiation therapy and chemotherapy. Therefore, it is important to consider the implications of therapy on the predisposition to flow stimulated cancer cell invasion. We qualitatively examined the expression of CXCR4 and CXCL12 in samples obtained from six patients diagnosed with glioblastoma prior to their receiving therapy and six patients after standard of care therapy (**Fig. 5A-H**). CXCL12 staining was generally more intense and widespread throughout the tissue than CXCR4, likely attributable to its role as a soluble cytokine. Pre-therapy samples were lightly positive for CXCR4 and many nuclei in the malignant regions did not appear to be associated with CXCR4 reactivity (**Fig. 5A-B**). Conversely, staining intensity appeared greater in samples obtained from patients who received therapy (**Fig. 5C-D**). Similar trends were also observed for CXCL12 staining, with perhaps a more dramatic difference between pre-therapy samples (**Fig. 5E-F**) and post-therapy samples (**Fig. 5G-H**).

**Figure 5:**
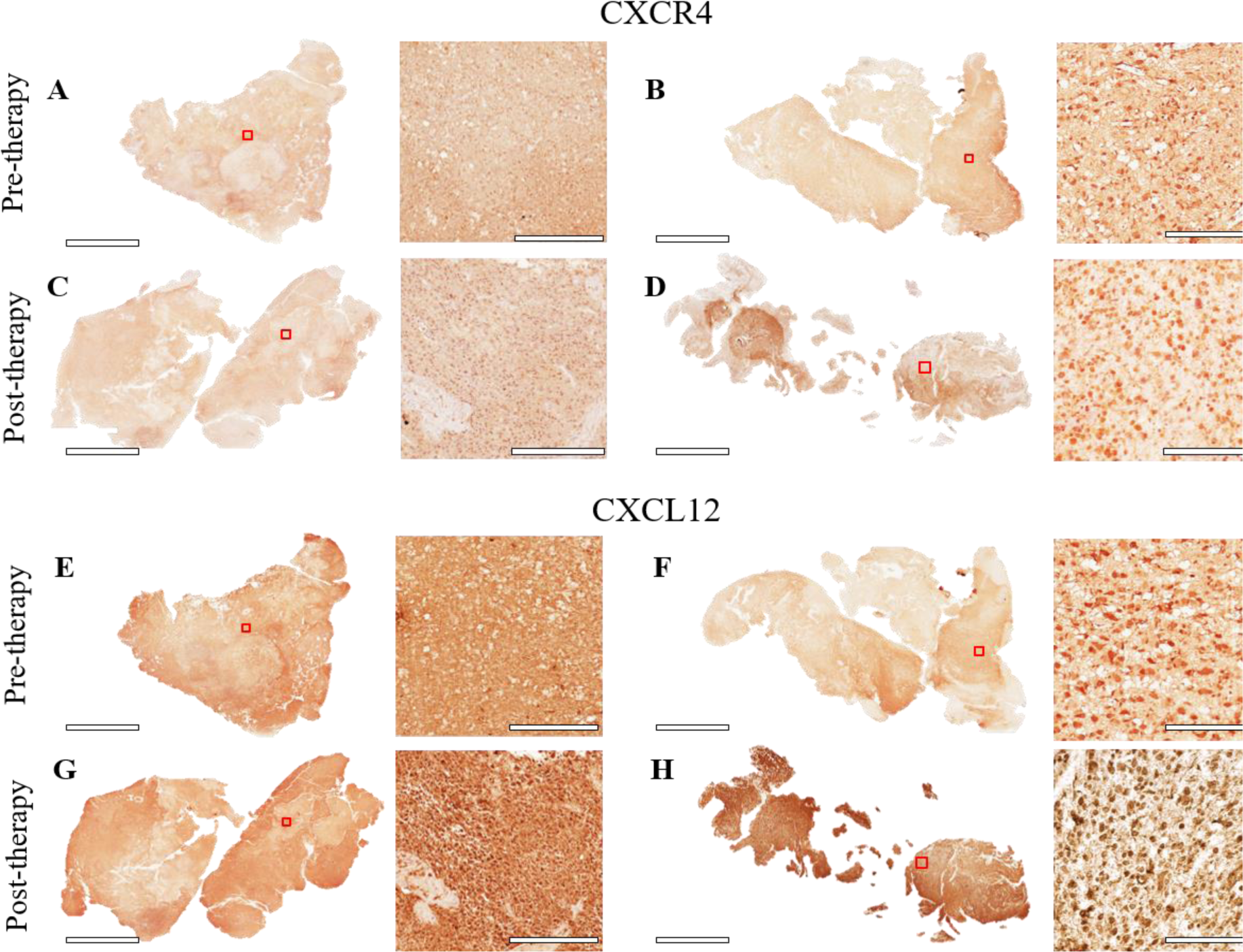
Cellular CXCR4 and CXCL12 –positivity increase in samples from patients who have received standard of care therapy. Twelve glioblastoma patient samples were grouped based on therapy status. Representative images are shown for four patients pre or post-standard of care temozolomide and radiation therapy immunostained for either CXCR4 (A-D) or its ligand CXCL12 (E-H) and counterstained with hematoxylin. The location of the inset image is outlined in red. Scale bars=7 mm for tissue scans, and 200 μm for inset images.

## Discussion

Interstitial fluid flow is a key component of normal physiology; however, emerging evidence suggests that this biomechanical force may also contribute to cancer malignancy (Munson and Shieh, 2014). The phenomenon is studied most extensively in breast cancer, where interstitial flow influences both the direction and magnitude of cancer cell migration and promotes activation of, and matrix remodeling by, relevant stromal cells^5,15-18^. Regarding GBM, paths of brain tumor dissemination correlate with bulk fluid pathways^19^. Only recently we showed that interstitial flow indeed increases invasion of both murine and human glioma cells through the chemokine receptor-ligand pair CXCR4 and CXCL12^6,7^. Nonetheless, the causal effects of flow on cancer cell invasion are currently known exclusively from *in vitro* experiments since previous *in vivo* data is only correlative. The goal of the current study was to elucidate the ability for interstitial fluid flow to directly stimulate glioma cell invasion in the brain.

The therapeutic technique of convection enhanced delivery (CED) was used to induce convective flow *in situ* within brain tumors, as evidenced by rapid contrast agent elimination. CED, a catheter-based method to by-pass transport limitations between the vasculature and the high-pressure tumor bulk, has been used experimentally and tested clinically for enhancing local perfusion of chemotherapeutics or other drugs in the treatment of GBM^10,20,21,22^. It was found that applying CED at 1 μL/min – the lower end of the clinically-relevant range of 1-5 μL/min^9^ – significantly increased GL261 cell invasion compared to untreated controls, based on analysis two days after flow application. This greater than 2-fold differential was more pronounced than in comparative *in vitro* experiments, suggesting an enhanced contribution of flow-stimulated invasion *in vivo.* One possible explanation for this pronounced effect is that CXCR4 expression was found to dramatically increase on GL261 cells upon implantation (from 8.13 ± 1.71% in 3D to 65.4 ± 5.19% *in vivo,* p<0.0001).

Because glioma cells can also express CXCL12, the ligand for CXCR4, it has been proposed that interstitial flow stimulates migration through a mechanism termed autologous chemotaxis^1,5^. Essentially, *in vitro* and *in silico* experiments suggest that fluid flow creates an anisotropic ligand gradient around individual cells in the direction of flow to stimulate directional migration. We previously showed using an agent based model that only small populations of CXCR4-and CXCL12-expressing cells are required to exhibit a flow response through this mechanism^7^. A drastic increase in CXCR4 expression *in vivo* may therefore greatly increase the likelihood of flow-stimulated invasion. Furthermore, although CXCL12 expression did not vary significantly between 3D culture and *in vivo,* CXCL12 is produced by other cells such as endothelial cells and astrocytes and is also present in the blood^23-25^. Therefore, ligand availability likely increases upon implantation, independent of cancer cell expression.

In the absence of any treatment, we observed moderate reactivity for CXCR4 phosphorylation, a known marker of receptor activation and signaling. Convective flow increased phospho-CXCR4 immunoreactivity both in the tumor and healthy brain tissue, suggesting the technique of CED may increase chemokine signaling and thus tumor cell dissemination. Additionally, there may be further implications of increased CXCR4 phosphorylation in the brain since activation of CXCR4 in glia can lead to increased neurotoxicity and pro-tumor phenotypes^26,27^. Additional studies are required to examine if the negative implications of CED (increased invasion and CXCR4 phosphorylation) are counter-balanced by the cytotoxic effects of an infused drug.

Other mechanisms have also been implicated in cancer cell response to flow. In particular, the hyaluronan-rich glycocalyx can mediate flow-driven mechanotransduction in part through the hyaluronan receptor CD44^28^. We observed here that CXCR4 signaling may be a primary mechanism by which the GL261 cell line responds to flow, but patient-derived glioma stem cells display heterogeneity in their dependence on CXCR4, CD44, or both for flow-stimulated invasion^7^. Additionally, blocking CXCR4 did not eliminate invasion entirely under static or flow conditions. Cancer cell invasion is a multifaceted process regulated by many mechanisms, as previously reviewed by Sayegh et al.^29^. Thus, here we identified CXCR4 as a regulator of flow-mediated glioma cell invasion, but other mechanisms can concurrently enhance infiltration into the brain.

While not examined here, it is important to consider that CED is most often used experimentally after standard radiation and chemotherapy. Previous work demonstrated that radiation induces tumor invasiveness by increasing tumor-derived CXCL12 at the invasive tumor border, which may enhance the potential for CXCR4 signaling^30^. Furthermore, irradiation of GL261 cells increased CXCR4 expression in a dose-dependent manner^31^. Using patient samples, we observed that post-therapy samples had increased immunoreactivity for both CXCR4 and CXCL12 compared to samples obtained pre-therapy. These observations suggest that the increased expression found in mice after therapy may hold true in humans. Beyond chemokine signaling, CXCR4 is also a purported marker of glioma stem cells^32^; therefore, increases to CXCR4 expression due to radiation may not only increase the potential for flow-stimulated invasion but also increase malignancy via cancer stem cell expansion. Our data imply that therapeutic use of CED, while advantageous for increasing drug transport and overall patient survival, may benefit from supplementation with CXCR4 blockade to preserve the benefit on drug permeance while preventing undesirable increases to cancer cell invasion.

## Materials and Methods

### In vitro invasion assays

GL261 invasion was assessed *in vitro* using 12-well (Millipore PI8P01250) or 96-well (Corning 3374) tissue culture inserts^1^. Cancer cells were seeded at 1 × 10^6^ cells/mL in 3D hydrogels, as described above. After 20 minutes of gelation, 15 μL of fresh medium was applied on top of the gels. Flow was initiated three hours later using serum-free medium, and cultures were maintained overnight. AMD3100 was used at 10 μM (Sigma A5602) to block the receptor CXCR4 while an excess of 100 nM CXCL12 (Peprotech 300-28A) was added to prevent chemokine gradient formation. The membranes were then fixed in 4% paraformaldehyde and counterstained using DAPI (Thermo Fisher D1306). An EVOS FL fluorescence microscope was used to acquire 20X images of the porous membrane bottom at five random locations for each sample^33^. The number of invading cells was manually counted for each technical replicate for n ≥ 4 biological replicates.

### Lentiviral transfection and in vivo tumor model

All animal procedures were approved by the Institutional Animal Care and Use Committees at the University of Virginia and Virginia Polytechnic Institute and State University. Lentivirus conferring expression of green fluorescent protein (GFP) under puromycin antibiotic selection was a generous gift from the laboratory of Dr. Kevin Janes. Murine GL261 were serially transfected with GFP lentivirus and purified by selection with 2 μg/mL puromycin (Thermo Fisher A1113803). For *in vivo* tumor studies, a burr hole was drilled into the skull of anesthetized C57BL/6 mice (5-8 weeks; Harlan Laboratories) at coordinates -2, +2, −2.2 (AP, ML, DV) from bregma. 100,000 GFP+ GL261 cells were inoculated in 5 μL at 1 μL/min, and the bur hole was sealed with bone wax. Ketoprofen was administered at 2 mg/kg for 48 hours to manage pain. One week later, the inoculation site was re-exposed, and a blunt-end 26 gauge needle was used to infuse 10 μL of 1 mg/mL biotinylated dextran amine at 1 μL/min. Ketoprofen was again administered at 2 mg/kg for 48 hours to manage pain.

### Flow cytometry

Triplicate wells of 100,000 GL261 cells were cultured in serum-containing medium overnight, either on 2D tissue culture plastic or in 3D hydrogels comprising 1.5% rat tail collagen (Corning 354236), 0.2% thiolated hyaluronic acid (Glycosil®; ESI Bio GS220), and 0.1% PEGDA (ESI Bio GS3006). The following day, cells were cultured with 10 μM Brefeldin A for 6 hours, harvested, pooled, and subjected to antibody labeling^7^. To assess expression *in vivo,* mice were inoculated with GFP+ tumor cells as above, and 14 days post-implantation mice were treated with 0.25 mg Brefeldin A for 6 hours via intraperitoneal injection^34^. The brains were then dissociated for analysis. Briefly, the ipsilateral cortical hemisphere was isolated into HBSS and slightly trimmed to reduce the number of non-cancerous cells. The tissue was minced using a scalpel blade, incubated in 5 mL of ACK RBC lysis buffer for 3-5 minutes at room temperature, and centrifuged at 1100 rpm for 5 minutes. An approximately equal volume of 1.5 mg/mL Liberase DL (Sigma 5466202001) was then added to digest the tissue for 30 minutes on a rocker at 37°C, pipetting up and down to ensure complete digestion.

The tissue slurry was then strained through a 40 micron cell strainer followed by 35 mL of HBSS. This solution was centrifuged at 1100 rpm for 5 minutes, and the isolated cells were resuspended and counted for flow cytometry. Primary-conjugated antibodies were used to stain for CXCR4 (eBiosciences 17-9991-80) and CXCL12 (R&D IC350C), along with appropriate isotype controls. Dead cells were stained using LIVE/DEAD^®^ Fixable Green Dead cell stain kit (Thermo Fisher L23101). Stained samples were run on a Millipore Guava flow cytometer for a minimum of 50,000 events, and the data was analyzed using Incyte software. A flow chart of the gating strategy is shown in **Supplemental Figure 1**. For data analysis, plots were gated based on data from single stained controls. *In vivo* samples were further gated on GFP+ cells to assess only GL261. All numbers are shown as percent of live, single cells.

### Magnetic resonance imaging

Animals were anesthetized and placed in a 7T Clinscan system (Bruker, Ettlingen, Germany) equipped with a 30-mm head coil. A T2-weighted image was taken through the head with the following parameters: repetition time (TR) = 5500 ms, echo time (TE) = 65 ms, field of view (FOV) = 20 mm × 20 mm with a 192 × 192 matrix, slice thickness = 0.5 mm, number of slices = 30, two averages per phase-encode step requiring a total acquisition time of about 5 min per mouse. For T1-weighted MRI, a 33-Gauge, blunt-end catheter was placed into the same coordinates for tumor implantation, and 10 μL of 25 mg/mL Glowing Galbumin (BioPAL Inc.) was infused at a rate of 1 μL/min. Approximately 30 minutes later, T1 images were acquired approximately 30 minutes, 1 hour, and 24 hours post-infusion according to the following parameters: TR = 500 ms, TE = 11 ms, FOV = 20 mm × 20 mm with a 192 × 192 matrix, slice thickness = 0.7 mm, number of slices = 22, two averages per phase-encode step requiring a total acquisition time of about 3 min per mouse. Contrast-enhanced T1-weighted images at time t=0 were subtracted from images at t=30 minutes to generate a difference heat map and visualize changes in contrast intensity over time.

### Tissue harvest and immunohistochemistry

Two days after convection enhanced delivery, tumor-bearing mice were overdosed on Euthasol solution and intracardially perfused with phosphate buffered saline (PBS). Brain tissue was quickly harvested and bisected at the center of the injection site. The brains were fixed overnight in 4% paraformaldehyde, cryopreserved in 30% sucrose, and sectioned at 12 μm using a Leica 1950 cryostat. Tissue sections were blocked in 3% serum and 0.03% Triton X-100 in PBS for 1 hour, then were incubated overnight at 4 °C with rabbit anti-pCXCR4 (Abcam ab74012) diluted in blocker buffer. The samples were washed three times with PBS and incubated for 1 hour at room temperature with goat anti-rabbit 660 diluted in blocking buffer. After washing again, the nuclei were counterstained using DAPI (Thermo Fisher).

### In vivo invasion quantification

Fluorescently labeled sections were imaged using an EVOS FL microscope. Five images were randomly taken around the tumor periphery for each of three sections 120 μm apart for each animal. The tumor border was identified based on GFP+ GL261 and nuclear staining, and a blinded investigator counted the number of GFP+ tumor cells beyond the border for each image. Data is presented as the number of invading cells per mm^2^ of tissue.

### Patient sample collection and immunohistochemistry

De-identified patient samples of glioblastoma were collected in accordance with the University of Virginia Institutional Review Board through the Biorepository and Tissue Research Facility with assistance from pathologists. Eight micron sections were deparaffinized in xylene and graded washed with ethanol:water to achieve rehydration. The samples were then subjected to boiling in citrate buffer for 30 minutes for antigen retrieval. The same primary antibodies were used as above: CXCR4 (Sigma GW21075) and CXCL12 (Abcam ab18919). Samples for CXCR4 were treated with goat anti-chicken IgY HRP secondary (Abcam ab97135) and those for CXCL12 with ImmPRESS™ horse anti-rabbit IgG HRP (Vector Labs MP-7401) prior to development with DAB. Images were acquired using an Aperio Slide Scanner and processed using ImageScope. Qualitative assessment was conducted on samples from a six patients per stain for each therapy status.

### Statistics

Analysis of Variance (ANOVA) was performed for comparisons of more than two groups, using a significance level of 0.05. If significance was identified within the dataset, t-tests were performed to determine significance between individual groups. Ratio paired t-tests were used to analyze all *in vitro* data; unpaired t-tests were used to compare *in vitro* data to *in vivo* flow cytometry data; and unpaired student’s t-tests were used to compare experimental groups for *in vivo* invasion. All graphed and reported descriptive statistics in the text are presented as mean ± standard error of the mean, unless otherwise stated. Inferential statistics are reported as statistics (degrees of freedom)=value, n per group, p value so that effect size can be determined from our reported data.

## Acknowledgments

The authors acknowledge Jack Roy and Stuart Berr in the University of Virginia Molecular Imaging Core, Dr. Frederick Epstein and Sophia Cui for MRI assistance, Dr. Scott Verbridge for manuscript feedback, and assistance from the University of Virginia Cardiovascular Research Center Histology Core and the Biospecimen and Tissue Repository Facility. Funding to JMM through ACS-IRG-81-001-29 and R01CA222563.

### Competing interests

The authors declare no competing interests.

### Authors’ Contributions

RCC contributed to study design, conducted data acquisition, analysis, and interpretation, and prepared the manuscript. CEB contributed to data acquisition for *in vitro* invasion. KMK conducted magnetic resonance imaging and generated difference maps. JMM conceived the study and supervised experimental design, analysis and interpretation of data, and manuscript preparation. All authors reviewed and edited the manuscript.

